# Microfluidic System for On-Demand Imaging of Organotypic Slices

**DOI:** 10.64898/2026.06.18.733098

**Authors:** Jana B. Petr, Emma Ferrari, Lorenz Hagelüken, Marco Cassano, Andreas Hierlemann, Mario M. Modena

## Abstract

Organotypic brain slice cultures (OSCs) provide a physiologically relevant model for recapitulating complex tissues in vitro. However, the longitudinal monitoring of dynamic processes in OSCs through high-resolution imaging is limited in standard culturing methods, as they rely on slice placement on porous membranes at the air-liquid interface to preserve tissue viability. Here, a new experimental paradigm based on a modular slice culture insert is introduced, which is compatible with static and perfused culturing, and facilitates longitudinal observation of cellular and tissue dynamics at high spatiotemporal resolution. Besides compatibility with conventional culturing in well plates, the insert is integrated into an automated microfluidic instrument to provide user-friendly operation while enabling precise temporal fluid control and compound delivery. The system is applied to monitor calcium responses and cellular morphology of virally labeled cerebellar astrocytes and Purkinje cells exposed to different concentrations of glutamate. Cellular responses were evaluated at baseline, during glutamate exposure, and 48 h and 96 h after stimulation to identify acute and long-term excitotoxic effects. Our results indicate that the developed system enables high-resolution monitoring of cellular morphology and dynamic events of the same intact tissue over multiple days, which provides a robust foundation for mechanistic investigations of neural signaling.

## 1. Introduction

Despite great advances in engineering, establishing models of complex tissues, which faithfully replicate the defined cytoarchitecture and composition of, e.g., brain tissue in vitro, remains challenging. A key example includes the cerebellum, which contains more than half of all neurons of the brain -including Purkinje cells and Bergmann glia - and accommodates highly specialized circuitry of the central nervous system.^1^ Owing to this complexity, the cerebellum remains understudied in experimental neuroscience, despite increasing evidence that cerebellar disfunction may be linked to several pathological conditions, such as ataxia, autism spectrum disorder, and Alzheimer’s disease.^2–4^ In vitro studies are often conducted using two-dimensional neuronal cultures due to their accessibility and ease of use. Two-dimensional cultures, however, do not adequately replicate the complex Bergmann glia-Purkinje cell interactions defining much of the synaptic neurotransmitter dynamics within the cerebellum’s tripartite synapses.^5^ Organotypic slice cultures derived from rodent and -more recently -human tissue provide three-dimensional model systems to overcome this limitation. These cultures largely preserve the complex cytoarchitecture, extracellular matrix composition, and cytokine signaling of native tissue.^6^ Additionally, neuronal connectivity and functional circuits can be re-established after slicing, which enables long-term studies in a functional-tissue context.^6,7^ Typically, commercially available tissue-culture inserts are used for ex vivo culturing of brain slices. The brain slices are placed on porous membranes at the air-liquid interface (ALI) to provide high oxygenation levels to the excised, metabolically still highly active tissues and to support tissue viability.^8^ However, the geometry of standard tissue-culture inserts is incompatible with longitudinal high-resolution confocal imaging under sterile conditions. As a result, the investigation of processes, such as intracellular calcium dynamics in neural populations, is currently restricted to endpoint analyses, which precludes a long-term tracking of evolving biological responses within tissue preparations. We have recently presented a platform for multi-day, high-resolution imaging of organotypic slices.^9^ There, we demonstrated that perfusion culturing could be used to sustain tissue viability in closed chambers to enable optical access to the tissues. However, the requirement to remain on the microscope stage after tissue loading and the incompatibility with standard ALI culturing between imaging sessions limited its applicability for longitudinal studies at relevant throughput.

To overcome these limitations, we have now engineered a modular, plastic-based culture insert that enables static culturing of organotypic brain slices in commercial 6-well plates and high-resolution imaging under perfusion conditions through confocal microscopy. The insert is also compatible with commercially available automated tissue staining instruments, like e.g., the LabSat from Bio-Techne Spatial,^10–12^ which we modified to enable perfusion with oxygenated medium to support tissue viability and functionality during imaging sessions. Additionally, perfusion enabled a precise temporal control of compound delivery, which facilitated the real-time observation of acute pharmacological responses during imaging. Furthermore, we made adaptations to ensure aseptic transfer of our tissue-culture inserts between a standard incubator and the imaging setup.

We validated the functionality of the developed system in experiments including brain-slice tissue culturing and repeated short- and long-term imaging sessions. The system was then used to investigate the response of Purkinje cells and cerebellar astrocytes to glutamate stimulation. Glutamate is the most important excitatory neurotransmitter in the central nervous system, playing a key role in synaptic plasticity, neural communication and neuroprotection.^13–15^ Dysregulations of glutamatergic signaling can lead to excitotoxicity, a pathological process driven by excessive levels of synaptic glutamate that results in calcium overload and subsequent neuronal degeneration. Glutamate dysregulations have been shown to be involved in several neurological diseases, such as autism spectrum disorders or Alzheimer’s disease.^16,17^

We used our system to investigate excitotoxic dynamics relevant to cerebellar neurodegeneration by exposing murine cerebellar slices to different concentrations of glutamate. We monitored morphological adaptations and changes in calcium dynamics at baseline conditions and at multiple time points during 96 hours after glutamate dosage.

## 2. Results

### 2.1. Insert for static culturing of organotypic brain tissue slices and repeated high-resolution live-cell imaging

Conventional air-liquid-interface (ALI) culturing of tissue slices, albeit effective for tissue viability, (i) entails limited optical access due to light scattering through the porous membrane or closed-lid systems and (ii) requires long-working distance objectives, which are incompatible with high-resolution imaging under aseptic conditions. Longitudinal investigations of dynamic processes, such as calcium imaging of neural activity at high resolution in organotypic tissue slices, currently require the preparation of several replicates per time point, as imaging is, in most cases, an end-point procedure carried out by immersing the sample in perfused medium, which affects sample sterility.

To overcome the current limitations, we devised a system compatible with both regimes, namely standard ALI culturing and on-demand perfusion, which enabled high-resolution imaging and continuous observation under aseptic conditions. Tissue culturing and perfusion were carried out on an insert (here referred to as LabSat insert) fabricated in inert plastic, polymethyl methacrylate (PMMA), to which a hydrophilic PTFE membrane was attached. Up to three brain slices per insert could be transferred onto the membrane immediately after tissue sectioning, and ALI culturing was then performed in a standard cell-culture incubator. The handling of the samples for daily medium exchange followed the same procedure as with traditional tissue-culture inserts, consisting of transferring the insert to a well with fresh medium. To perform high-resolution imaging, the insert was first mounted on the imaging holder under aseptic conditions. Subsequently, the holder was moved to the confocal microscope and connected to the fluidic, pneumatic, and electronic components of the LabSat instrument. High-resolution imaging using immersion objectives could then be performed without risking contamination of the tissue slices. Slice health during imaging was ensured by perfusion with culture medium that had been equilibrated with carbogen (95% O2, 5% CO2) before delivery to the slices. At the end of the imaging session, the imaging holder was removed from the fluidic system, moved to a laminar flow hood, the insert was removed and returned to the 6-well plate, and maintained under ALI conditions in the incubator (**Figure 1A**). The design parameters of our insert were optimized for placement in a standard 6-well plate, using 1 ml of medium per well. In this aspect, we replicated the conditions of the current gold standard method based on commercially available tissue-culture inserts (**Figure 1B**). In contrast to standard inserts, our design featured a thin (500 µm) ring for membrane support to promote compatibility with high-resolution water immersion objectives and to prevent potential damage to the tissue slices during clamping for enabling perfusion (**Figure 1C**). This technological feature enabled longitudinal imaging assays across scales, from tissue level to subcellular details (**Figure 1D**). In parallel, we modified the LabSat instrument, originally devised for immunostaining of tissue sections, to be compatible with imaging of live samples and to accommodate our insert: (i) The sample holder was modified to host and align the LabSat insert with removable alignment structures made of polycarbonate, a temperature and chemical-resistant plastic material that is compatible with autoclaving and periodic washing with ethanol (**Figure S1**); (ii) the LabSat imaging holder was rendered detachable from the instrument to enable transfer of the insert from the culture plate to the holder in a laminar flow hood and prevent sample contamination; simple connections for the fluidic, pneumatic and electronic components enabled to quickly detach and reattach the holder at the microscope location (**Figure S2**); (iii) protocols controlling the flow (25-30 µl/min) and temperature of the PMMA insert were generated in the LabSat software and validated using an integrated inline flow sensor downstream of the imaging holder to support sample viability and automate fluidic control during the imaging assays (**Figure S3**).

**Figure 1:**
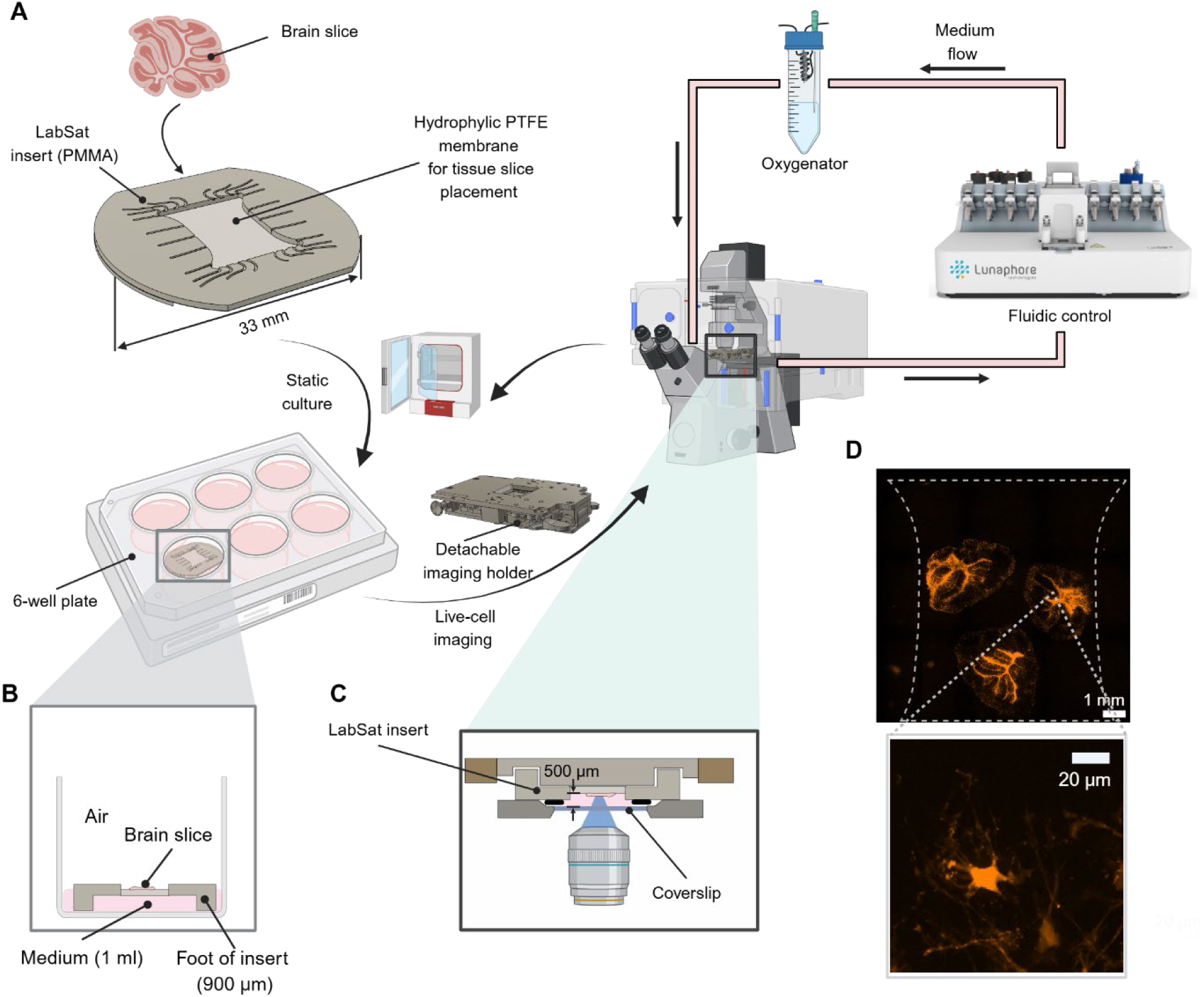
**A:** Design of the LabSat insert for organotypic slice culturing and workflow to integrate the insert into a commercially available microfluidic system for recurrent live-cell imaging. **B**: Cross-section of the insert placed in a 6-well plate. Foot height was chosen to work with 1 ml of medium, as in standard culturing protocols. **C**: The close-up shows the cross section of the holder with the loaded insert using an inverted microscope with a working distance of 500 µm: **D:** The system allows for visualizing events from tissue level to single-cell level. The dotted lines indicate the insert borders. Orange: GFAP-driven mCherry.

### 2.2. Static on-insert culture supports tissue recovery and Purkinje cell layer reorganization

Reorganization and survival of neuronal networks in organotypic slices after tissue sectioning require tightly regulated culturing conditions to ensure sufficient delivery of oxygen and nutrients to the tissue. We first sought to confirm that our insert allowed for long-term static culturing and recovery of organotypic slices. To this end, we used the size of the Purkinje cell soma and the Purkinje cell density as metrics to confirm the health of the Purkinje cell layer during static culturing. After tissue slicing, cerebellar slices were transferred to commercial tissue culture inserts and to the LabSat insert, followed by 14 days of static culturing to allow for tissue recovery and reorganization. Slices were then fixed and stained using immunofluorescent neuronal and glial markers to assess the morphology of Purkinje and glial cells (**Figure 2A**). The Calbindin staining confirmed that the Purkinje layer reorganized into a continuous, uninterrupted cell layer, which is indicative of good tissue health (**Figure 2B**). High-magnification images of the cerebellar lobules confirmed correct cytoarchitectural organization, displaying layered and adjacent molecular layer, Purkinje cells, and granular cell layer, followed by distinct and organized axonal tracts (**Figure 2C**). Finally, quantification of Purkinje cell volume (**Figure 2D**) and Purkinje cell density (**Figure 2E**) did not show significant differences between our culturing insert (mean Purkinje cell volume of 1416 ±1212 µm^3^, mean density of 0.0027 ± 0.0006 cells/µm^2^) and standard tissue-culture inserts (mean Purkinje cell volume of 1459 ± 1029 µm^3^, mean density of 0.0020 cells ± 0.0005 cells/µm^2^) and were in close agreement with previously reported in vivo data.^18^ These results confirmed that the developed insert was suitable for culturing of organotypic slices.

**Figure 2:**
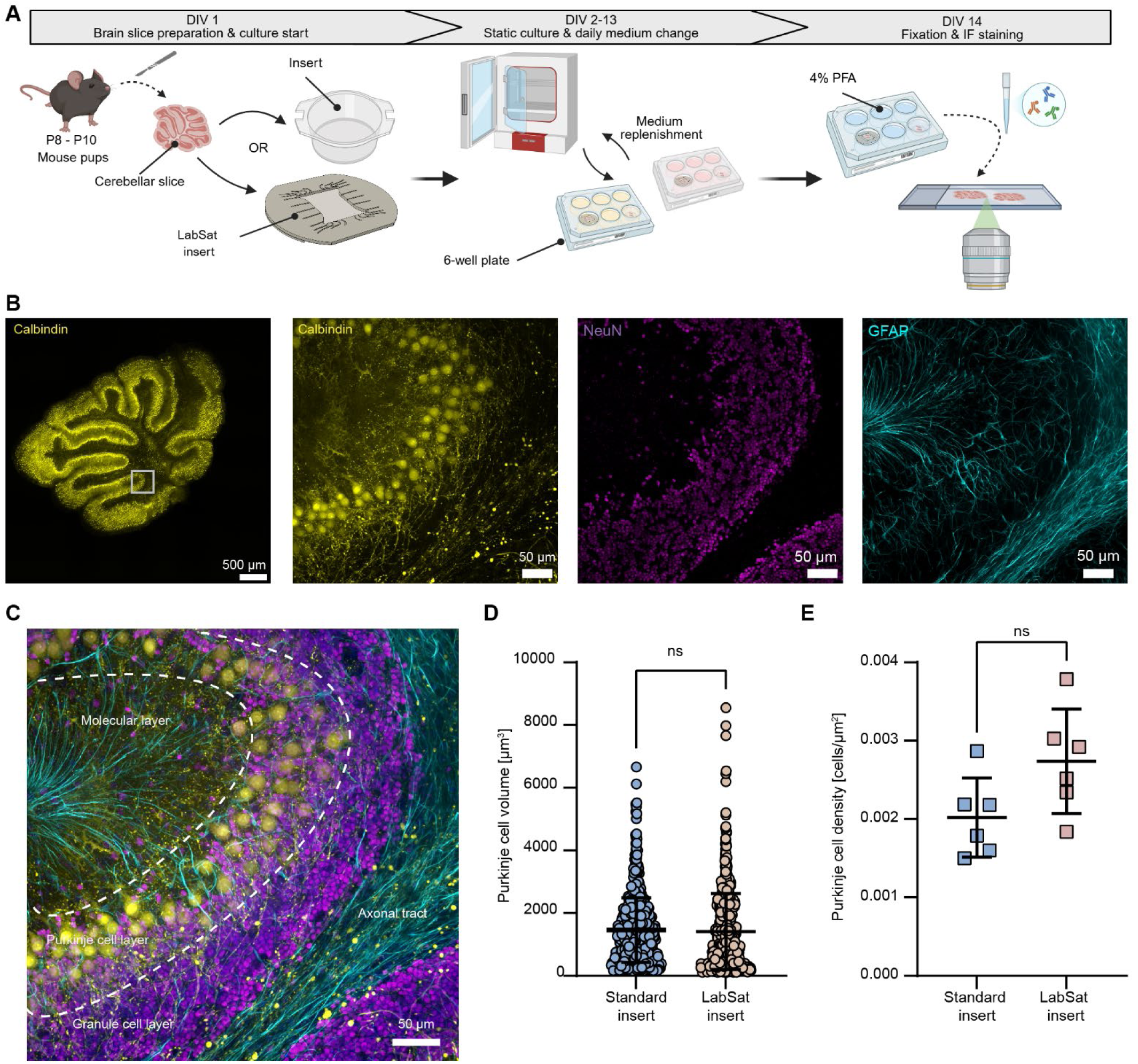
**A**: Experimental approach to assess the suitability of the developed insert for organotypic slice culturing. **B**: Left, representative image (maximum intensity projection) of a cerebellar slice cultured on the LabSat insert and showing a continuous Purkinje cell layer across the whole slice (yellow: anti-Calbindin); the three insets show higher-magnification images of the highlighted region to display the morphology and organization of the Purkinje cells (yellow: anti-Calbindin), the granular cell nuclei (magenta: NeuN) and the astrocytes (cyan: anti-GFAP). **C**: Composite image of insets in B showing the organization of the cerebellar slice after 14 days of static culturing. **D-E**: Comparison of Purkinje cell soma volumes and cell density between slices cultured on standard cell-culture inserts (blue) or on LabSat inserts (red). Welch’s t-test was used for statistical analysis of cell volume and cell density data.

### 2.3. Perfusion with oxygenated medium sustained tissue viability

Next, we sought to investigate whether our system supported brain tissue health during extended culturing under pulsated and oxygenated perfusion in a closed environment. We cultured cerebellar slices under static conditions at the ALI for 14 days in vitro. On day-in-vitro (DIV) 14, the insert with mature organotypic slices was transferred to the imaging holder, and slices were cultured under perfusion for 24 hours (**Figure 3A**). We evaluated baseline viability after the initiation of perfusion culture with a viability staining, which contained Hoechst to stain cell nuclei and a cleaved caspase-3/7 marker to identify apoptotic cells. The same staining procedure was then repeated after 24 hours under perfusion. We did not observe a pronounced presence of caspase-3/7^+^ cells at the beginning or after 24 hours of perfusion culturing (**Figure 3B**). To confirm that the cerebellar slices were viable and that the lack of apoptotic signals was not caused by tissue that was already necrotic at transfer to the imaging holder, we took advantage of the fluidic system to perfuse 40 µM doxorubicin for 2 hours and induce tissue apoptosis. After doxorubicin exposure, the slices displayed a high level of positive caspase signal, resulting in a 250-fold increase in caspase signal intensity with respect to baseline conditions and the 24-hour recordings (**Figure 3C**). This finding confirmed that slices were alive when they were loaded into the system and that perfusion culturing did not affect slice viability.

**Figure 3:**
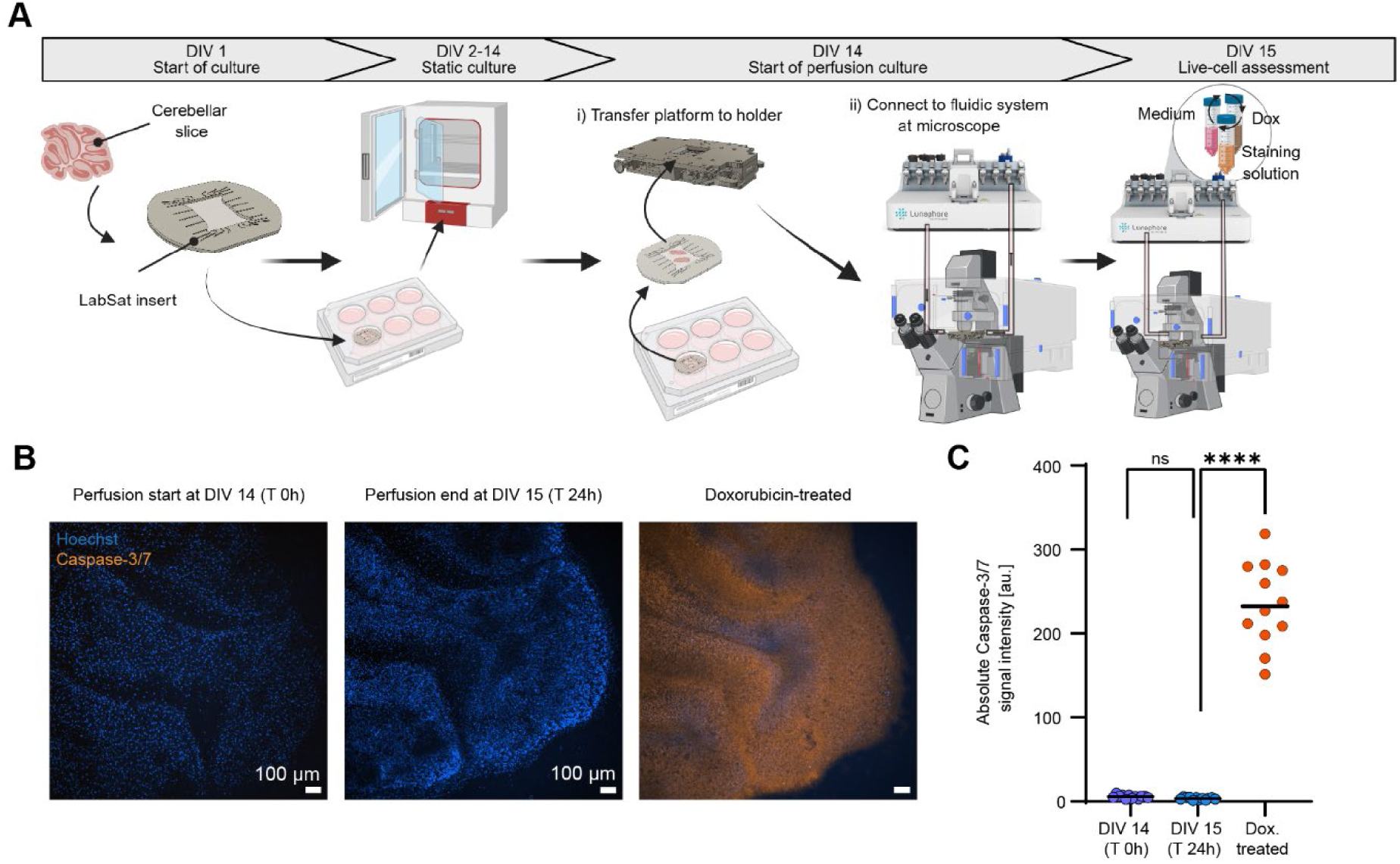
**A:** Schematic workflow of the experimental timeline to assess tissue viability under prolonged perfusion. **B**: Representative maximum intensity projections of the same ROI stained with Hoechst (blue) and cleaved Caspase-3/7 (orange) at the start of the perfusion culture, 24 h into the perfusion, and after 2 h treatment of 40 µM doxorubicin. **C**: Background-subtracted absolute Caspase-3/7 intensity at 0 h and 24 h of perfusion and after doxorubicin treatment. One-way ANOVA was used to derive statistical significance using n = 12 ROIs from 3 slices.

### 2.4. Acute and long-term effects of glutamate on Purkinje cell activity

We then used our system to investigate acute and long-term cytotoxic effects of excess glutamate exposure. Glutamate is an essential neurotransmitter, which, in the cerebellum, has an excitatory effect on Purkinje cells mediated by parallel fibers extending from the granular cells. Owing to the key role of glutamate signaling, disbalances in global glutamatergic levels are known to affect calcium dynamics and cell morphology.^19–22^

To specifically label Purkinje cells, we transduced the cerebellar slices at DIV 4 and DIV 6 with a GCaMP8m AAV under the expression of the L7-6 promoter. The slices were then cultured for a minimum of 8 additional days to support tissue reorganization and a stable expression of the GCaMP reporter before glutamate exposure. After tissue maturation, the inserts were transferred to the imaging holder, and baseline calcium dynamics were acquired. Subsequently, the slices were perfused with 0.1 and 1 mM glutamate solutions for 30 min using the fluidic system. After glutamate dosage, the samples were returned to standard medium conditions, and calcium dynamics were recorded for ∼1 hour. At the end of the imaging assay, the inserts were removed from the holder, transferred to a 6-well plate, and placed in an incubator until the next imaging sessions at 48 and 96 hours post glutamate stimulation (**Figure 4A**).

**Figure 4:**
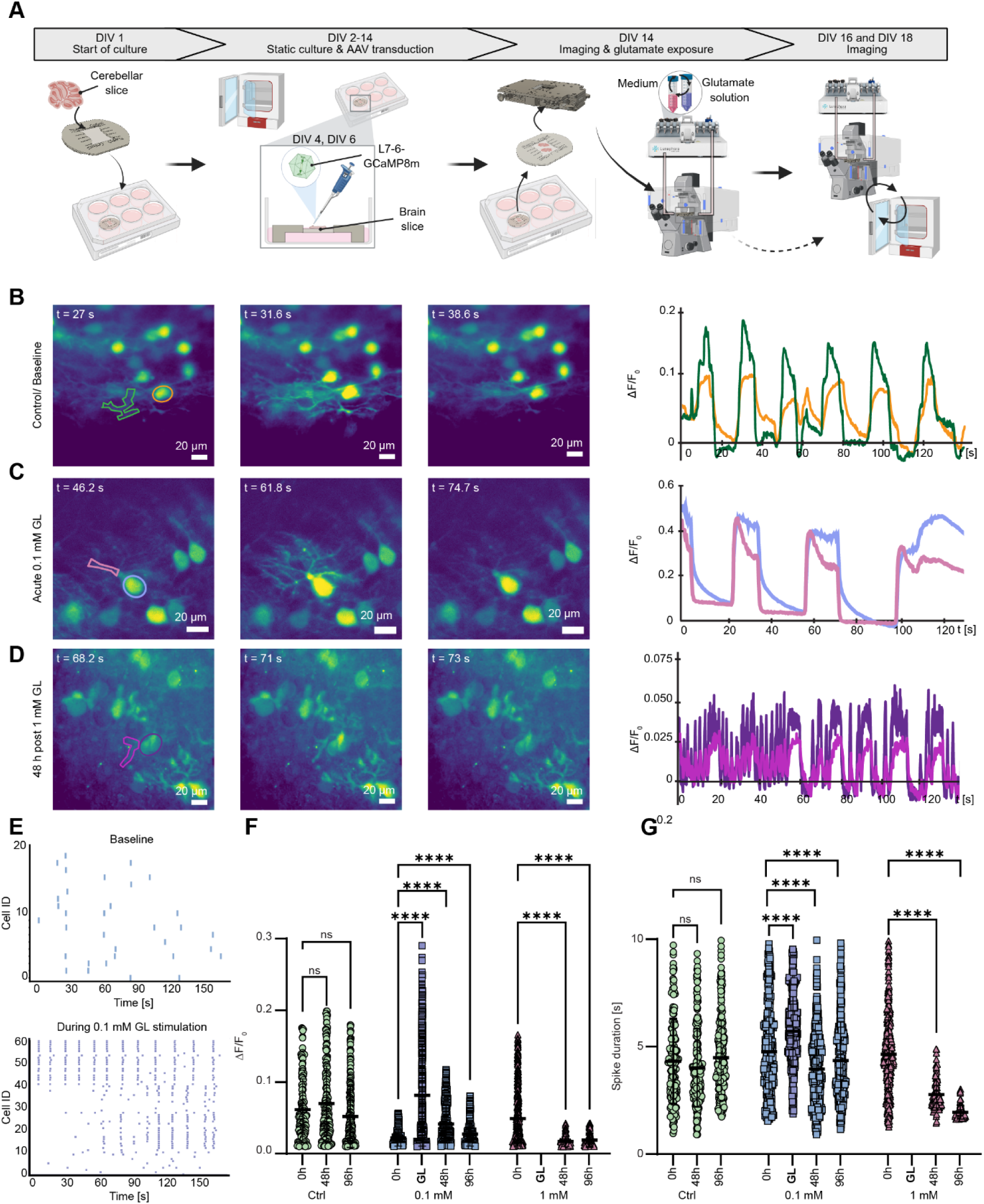
**A**: Experimental procedure to investigate Purkinje cell calcium dynamics after glutamate exposure. **B-D**: Time frames of representative calcium spikes in Purkinje cells prior to glutamate exposure, during 0.1 mM glutamate exposure and 48 h post 1 mM glutamate exposure. On the right, ΔF/F0 traces of calcium events in somas and processes visualized in B- D. **E:** Scatter plots showing active cells and the corresponding firing events before (top) and during (bottom) stimulation with 0.1 mM glutamate. **F-G**: Average maximum ΔF/F0 and spike duration per cell and condition prior to glutamate exposure (0 h), during glutamate exposure (GL), and 48 h and 96 h post exposure (n= 5 slices per condition). Empty columns indicate conditions where no spikes were detected. Statistical analysis was performed using a one-way, non-parametric ANOVA test.

The high temporal and spatial resolution attainable with our system enabled us to record calcium dynamics in both somas and fine branches of the dendritic trees of the Purkinje cells. Image acquisitions of unexposed Purkinje cells revealed rhythmic spiking activity with larger intensity changes observed in the dendritic tree than in cell somas, which corresponded to the trends observed in in vivo recordings of awake mice ^23^ (**Figure 4B)**. However, during acute exposure to 0.1 mM glutamate, dendritic structures and somas of Purkinje cells showed similarly large intensity changes (**Figure 4C, Supplementary Video 1**). Finally, during 1 mM glutamate exposure, we were unable to record any calcium dynamics (**Supplementary Video 2**): the cells exhibited a much higher resting signal level so that no dynamic signals were detected, potentially due to a saturation of the glutamate buffering in the Purkinje cells.^22^ Re-assessment 48 h after 1 mM glutamate exposure revealed a low number of active cells, whose firing pattern featured non-rhythmic spikes in both the dendritic trees and the somatic regions (**Figure 4D**).

Slices stimulated with 0.1 mM glutamate showed an increase in firing frequency and a general higher activity level, i.e., a larger number of active cells during acute exposure (**Figure 4E**). This increase in cell activity was also reflected in higher fluorescence intensity changes, which peaked during acute exposure and showed a consistent higher average value with respect to baseline conditions throughout the 96-hour observation period (**Figure 4F**). Purkinje cells stimulated with 1 mM glutamate, on the contrary, featured reduced spike amplitude at 48 and 96 hours post stimulation. Glutamate stimulation also led to alterations in spike duration, which increased during acute exposure of cells to 0.1 mM glutamate, before decreasing below baseline values at 48 and 96 hours after stimulation (**Figure 4G**). Spike duration decreased to a much lower value for Purkinje cells treated with 1 mM glutamate, which displayed calcium events that were more than 2-times shorter at late-time-point acquisitions. Control conditions did not cause any significant variation in fluorescence intensity dynamics and spike duration throughout the whole observation period, which confirmed that repeated loading and imaging of the sample did not have an effect on the calcium dynamics of the Purkinje cells.

### 2.5. Glutamate exposure alters Purkinje cell morphology

In parallel to calcium dynamics, we investigated how excess glutamate affected the morphology of Purkinje cells. To this end, we assessed the area of the cell soma, the dendrite thickness and the number of dendritic branches (**Figure 5A**, **Figure S4A-C**). Glutamate exposure resulted in both acute and long-term changes of cell morphology: no acute effect on the soma area was detected for 0.1 mM glutamate exposure, while cells displayed a ∼1.5-fold increase in soma area upon exposure to 1 mM glutamate (**Figure 5C**). A similar effect was also recorded for Purkinje cells exposed to even larger glutamate concentrations, namely 3 and 10 mM (**Figure S4D**). In contrast, at 48 hours post exposure, 0.1 mM glutamate resulted in soma enlargement, while the soma area reverted to the average baseline value for the 1 mM condition. Finally, at 96 hours post exposure, both conditions displayed a significant increase in cell soma. Due to the high cytotoxic effect of large glutamate concentrations, Purkinje cells exposed to 3 and 10 mM could not be monitored for the whole assay duration.

**Figure 5:**
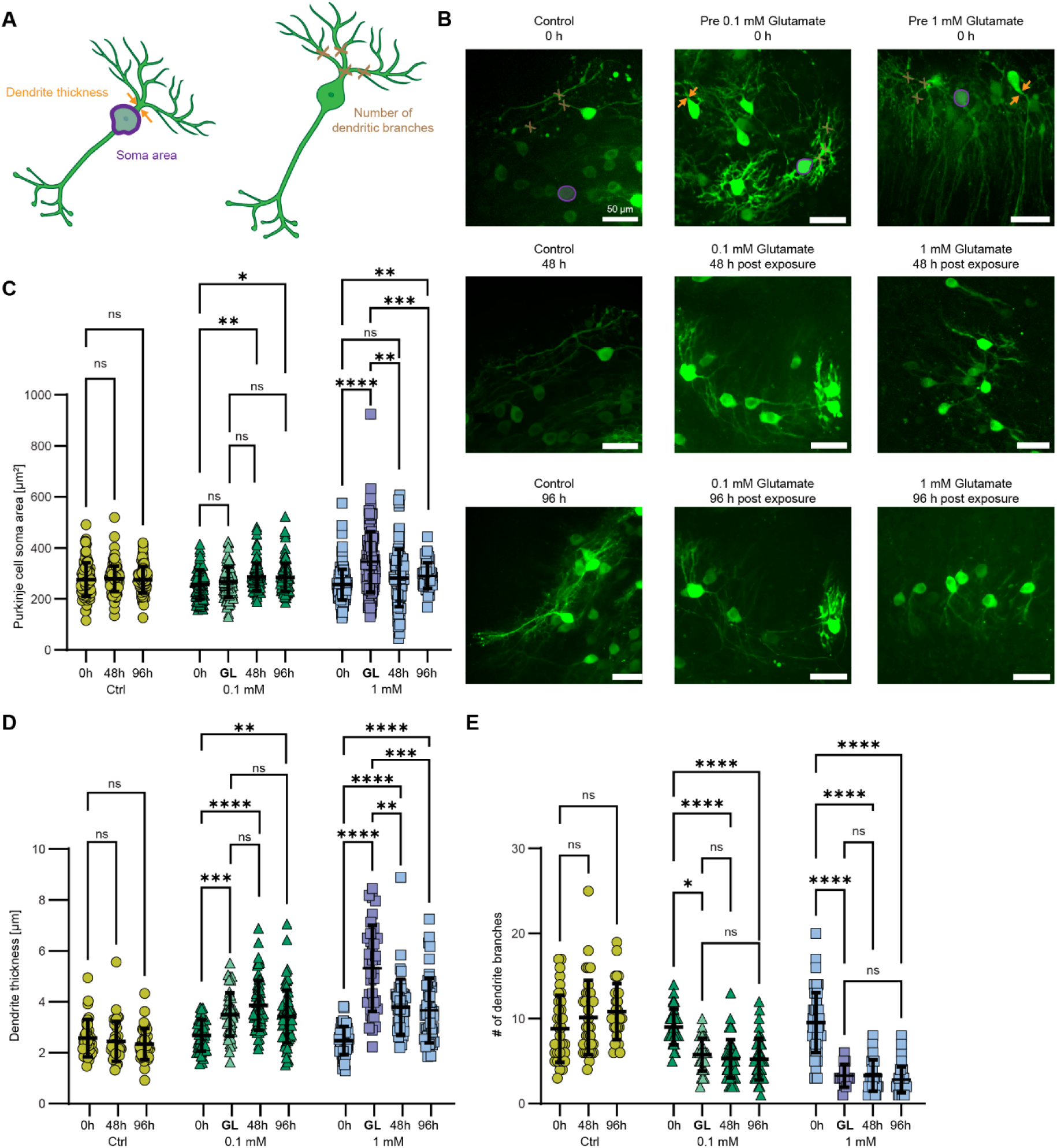
**A:** Cartoon showing the assessed morphological changes in Purkinje cells. **B:** Representative maximum-intensity-projection images of cells before, during, and after glutamate exposure. **C:** Area of Purkinje cell somas. **D:** Purkinje cell dendrite thicknesses. **E:** Number of dendritic branching points. Morphological parameters were acquired before glutamate exposure (0 h), during glutamate exposure (GL), and 48 h and 96 h post glutamate exposure. Data of n = 5 slices per condition. Statistical analysis was performed using a one-way, non-parametric ANOVA test.

Glutamate exposure also affected dendrite thickness and the number of dendritic branches (**Figure 5D-E and Figure S4E-F**), resulting in both a rapid and sustained increase of dendrite thickness and a corresponding reduction of the number of dendritic branches. The overall effect was more rapid and pronounced for the 1 mM condition, where dendrite thickness during stimulation was larger than at the 48-hour time point. Dendrite thickness then appeared to decrease from 48 to 96 hours for both conditions; however, dendrite thickness did not return to the baseline value within the experimental window. As already mentioned, a reduction in the number of dendritic branches was detected during stimulation, with a more pronounced effect for the 1 mM dosage in comparison to 0.1 mM, and remained stable throughout the whole assay duration, as no recovery was observed. Finally, we confirmed that perfusion alone did not affect dendrite thickness and the number of branches as evidenced by the lack of any significant change throughout the assay.

### 2.6 Acute and long-term effects of glutamate on astrocyte morphology and calcium dynamics

Astrocytes are the most abundant glial cell type in the cerebellum and play a key role in glutamatergic homeostasis. In the tripartite synapse, astrocytes are responsible for the uptake of excess glutamate in the synaptic cleft, for its transformation into glutamine, and the respective re-uptake into presynaptic vehicles.^24^ Consequently, high glutamate concentrations are expected to affect astrocytes, a mechanism that remains understudied in the cerebellar context.

Similar to the experimental workflow for investigating the response of Purkinje cells to glutamate stimulation, we used the static recovery period after slice preparation to transduce with and express the calcium sensor GCaMP6f and the mCherry protein under the hGFAP promoter (**Figure 6A**). Labeling the astrocytes with a mCherry fluorescent protein enabled us to more reliably visualize astrocyte morphology, regardless of their calcium dynamics.

**Figure 6:**
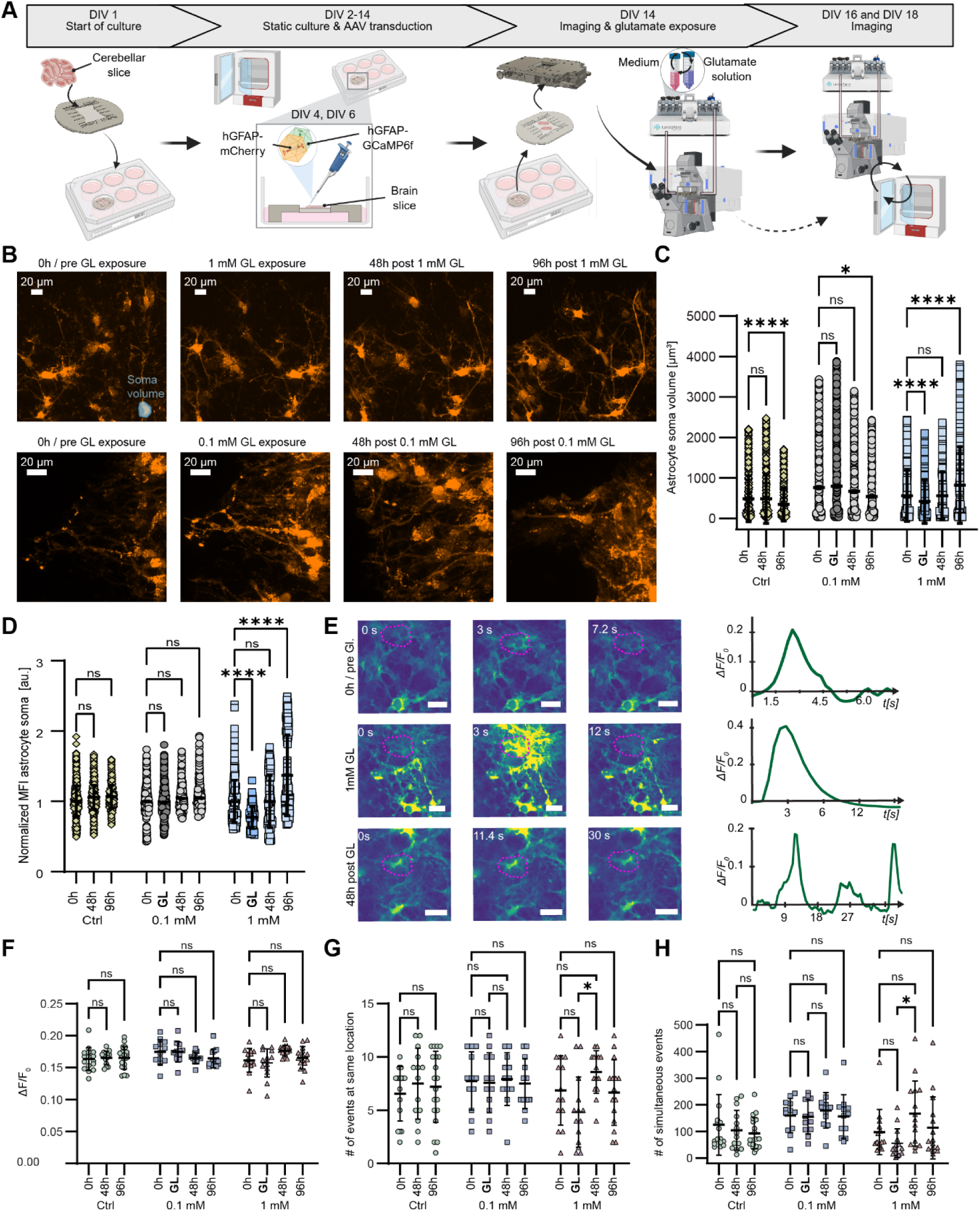
**A:** Experimental workflow to study morphological changes and calcium dynamics in cerebellar astrocytes upon glutamate exposure. **B**: Representative maximum intensity projections of astrocytes before (0 h), during (GL), and 48 h and 96 h post glutamate exposure applying 1 mM glutamate (top row) or 0.1 mM glutamate (bottom row). Orange: GFAP-driven mCherry. **C**: Astrocytic soma volume. **D**: Mean fluorescence intensity (MFI) of the soma volume. **E**: Time frames and fluorescence traces of a calcium event before (0 h), during (GL), and 48 h after 1 mM glutamate exposure. Maximum fluorescence intensity variations ΔF/F0 per ROI. **G**: Number of events at the same location per ROI. **H:** Number of simultaneously occurring events. In **F**, **G** and **H**, each point indicates the median value per ROI. Data was acquired from n = 3-5 slices per condition. Statistical analysis was performed using a non-parametric, one-way ANOVA test.

In line with longitudinal observations of the Purkinje cells, we assessed morphological parameters, including astrocyte soma volume and mean fluorescence intensity (MFI) of the soma volume, and the calcium dynamics of astrocytes in the slices before and during dosage, and 48 and 96 hours after glutamate stimulation.

Morphological measurements were performed by detecting the expression of the fluorescent mCherry protein (**Figure 6B**). Astrocytes stimulated with 1 mM glutamate showed an initial decrease in soma volume during exposure and recovered to baseline values 48 hours after exposure. Astrocyte soma volume continued to increase until the next time point at 96 hours, where the measured soma values significantly exceeded baseline values. In contrast, we did not observe any change in astrocyte soma volume in cells treated with 0.1 mM glutamate during exposure and 48 hours post exposure. However, we observed a decrease in soma volume at the 96-hour timepoint for slices stimulated with 0.1 mM glutamate and for control slices (**Figure 6C**). The reduction in astrocytic soma volume at 96 hours in control slices may be a result of some adaptation of the astrocytes to perfusion, while astrocytes exposed to the highest glutamate concentration showed an opposite effect, with continuously increase in soma size after glutamate exposure.

The expression of mCherry, and, therefore, its mean fluorescence intensity (MFI), is directly linked to GFAP expression, the level of which is an indicator of astrocyte reactivity. Therefore, we evaluated the MFI of mCherry in the astrocyte soma as a proxy for astrocyte reactivity (**Figure 6D**). Astrocytes under control medium conditions did not show any increase in MFI during the assay, which indicates that perfusion did not affect mCherry expression. Similarly, we did not detect any variation in mCherry MFI for slices exposed to 0.1 mM glutamate. In contrast, the MFI of cells exposed to 1 mM glutamate first decreased during acute acquisitions, before showing a continuous increase throughout the 48- and 96-hour timepoint observations, finally exceeding the baseline value. The increase in mCherry MFI likely indicates an increase in the reactivity of the astrocytes exposed to the highest glutamate concentrations.^25,26^

We also evaluated the variations in calcium dynamics caused by glutamate exposure. Owing to the alignment features of the chip in the imaging holder, we were able to assess the effects on the same cells across multiple time points and to extract the related fluorescence traces (**Figure 6E**, **Supplementary video 3-4**). Glutamate exposure appeared to influence the overall dynamics of astrocyte calcium signaling. However, upon performing a quantitative analysis of calcium imaging metrics, namely the maximum fluorescence intensity variations, the number of events at a certain location during acquisition, and the number of simultaneously active cells, we were not able to detect any statistically significant difference, neither during acute exposure, nor as a long-term response to glutamate exposure (**Figure 6F-H**). Astrocytes exposed to 1 mM glutamate showed a decreasing trend in the number of events at the same cell location and in simultaneously occurring events, possibly due to a temporary metabotropic receptor desensitization,^27^ followed by a statistically significant increase at 48 hours. Stimulation with 0.1 mM glutamate did not alter calcium dynamics with respect to any of the assessed parameters. Intracellular astrocytic calcium dynamics in untreated slices remained unchanged throughout the experiment timeline, indicating that the alterations between static ALI and perfusion culture did not seem to affect calcium dynamics.

## 4. Discussion

In this study, we present the development of a culture insert for organotypic tissue slices, its use with standard well plates, and its integration into a commercial fluidic system for on-demand imaging of dynamic biological processes. The static configuration allowed us to replicate the standard workflows for organotypic slice culturing on tissue-culture inserts. During static culturing, the slices recovered from the lesions caused by sectioning^28^ and were subjected to further experimental procedures, such as viral transduction. After this recovery period, we performed a perfused imaging assay by transferring the insert to the imaging holder, and acquired fluorescence images of unstimulated and glutamate-stimulated slice cultures to assess cell morphology and calcium dynamics until up to 96 hours after stimulation. Between imaging sessions, the platforms were returned to ALI culturing in a standard CO2 incubator.

The modularity of the experimental design and setup enables the use of the insert and the corresponding overall system for longitudinal studies, supporting the recurrent imaging of dynamic processes in a tissue context. For example, organotypic slices from various brain regions, including human tissues, have been maintained at the ALI for applications such as cancer invasion studies,^7,28–32^ and our system would allow to monitor cellular invasion and response to drug treatment during multi-day assays. Since the static operation of our insert is identical to conventional culturing, established protocols for existing models can be transferred without modification. To confirm that our insert does not affect tissue viability and recovery after slicing, we evaluated the reorganization of Purkinje cell layers after slicing, which yielded the same results that have been obtained with standard inserts.

Additionally, we demonstrated that our system is compatible with prolonged imaging sessions without affecting slice viability. The slices were periodically perfused with oxygenated medium for 24 hours, whereas no increase in the number of apoptotic cells was detected between the start and the end of the imaging session.

We then applied our system to investigate the response of cerebellar slices to high glutamate concentrations. We first monitored intracellular calcium dynamics in Purkinje cell somas and dendritic trees. Under standard medium conditions, fluorescence traces of soma and dendritic trees of the same cell resembled in vivo calcium dynamics.^3,33,34^ Upon exposure to the lowest tested glutamate concentration (0.1 mM), we observed rapid and irreversible alterations of calcium activity in Purkinje cells. Dosage of 1 mM glutamate then appeared to overstimulate Purkinje cells, which resulted in very low signal-to-noise ratios and no detectable neuronal activity. This effect likely reflects calcium-mediated excitotoxic silencing, whereby excessive receptor activation leads to transient loss of electrophysiological outputs.^35^ As our system enabled to re-image the same Purkinje cells under baseline conditions, during glutamate exposure, as well as 48 hours and 96 hours after stimulation, we followed the response development of the Purkinje cells throughout a 4-day assay. 48 h post exposure, the fluorescence traces of cells stimulated with 1 mM glutamate showed a drastic variation in signal dynamics and periodicity, which was reflected in decreased spike duration and decreased intensity changes in the pooled data. Purkinje cells treated with 0.1 mM glutamate featured acutely increased spiking activity upon exposure, which is consistent with the reported glutamate-mediated opening of NMDA and AMPA receptors of Purkinje cells.^36,37^

Besides the observed alterations in calcium dynamics, we observed glutamate-induced dose-dependent morphological changes consistent with excitotoxic swelling. The morphological changes were partially reversible over the 96-hour observation window, but the parameters remained different from baseline values. Our measurements are in concordance with reported observations of morphological changes in Purkinje cells in neurological diseases linked to alterations in glutamate homeostasis.^38,39^

Analogous to Purkinje cells, we also measured the response of astrocytes to the same glutamate dosages. Unlike Purkinje cells, astrocytes did not show significant changes in calcium-signaling fluorescence intensity during glutamate exposure, which could be related to their role of recycling and removing excess glutamate, as well as their ability to regulate and buffer intracellular calcium.^40^ We observed a trend towards an acute decrease in the number of astrocytic calcium events upon 1 mM glutamate exposure, which may reflect mGluR5 desensitization. This is a well-characterized mechanism, whereby sustained high-dose receptor activation leads to receptor uncoupling and attenuation of downstream calcium signaling.^27^ Although the differences between the applied conditions did not reach statistical significance at the slice level, likely due to inter-slice variability,^41^ we identified a statistically significant increase in calcium event frequency from the dosage timepoint to 48 h post-exposure. This delayed upregulation is in agreement with the onset of reactive astrogliosis, during which reactive astrocytes are known to amplify Ca²⁺ wave propagation and increase gliotransmitter release as part of a neuroprotective response.^42^ This event interpretation is supported by the concurrent increase in GFAP expression and in soma volumes at 48 and 96 hours, two phenomena indicative of astrocyte reactivity.^25,26^ The MFI of astrocytic soma mCherry intensity initially decreased upon 1 mM glutamate exposure, which could be caused by a transient dysfunction of GFAP expression due to calcium overload, as previously described in literature.^43^ The following gradual increase in MFI in combination with increases in soma volumes could point towards an increased reactivity of the astrocytes with upregulation of GFAP as a neuroprotective mechanism.^44^

In summary, the described case studies confirm the suitability of our culture insert and fluidic system for longitudinal studies of dynamic neural processes in organotypic brain slices. We demonstrated that our microfluidic system enables stable, longitudinal imaging over multiple days without compromising tissue viability and functionality. Applied to glutamate excitotoxicity, the system revealed a vulnerability difference between Purkinje cells and astrocytes and provided dose-dependent and time-resolved signatures. Although the current configuration accommodates only one insert for imaging at a time, up to three slices per insert can be cultured and imaged simultaneously, which partially compensates for this constraint. Additionally, inserts can be rapidly loaded and removed from the holder, thereby enabling the analysis of multiple conditions per day.

Finally, the results presented here were acquired using murine cerebellar slices. However, the system’s compatibility with established ALI protocols renders it a broadly applicable tool for longitudinal studies across brain regions and disease contexts also for human tissue.

## 3. Experimental Section

### 3.1. LabSat insert fabrication

The design of the LabSat insert was made in Fusion 360 (Autodesk Inc., San Francisco, CA, United States). The insert featured an outer diameter of 33 mm and was manufactured by micro-CNC milling (Datron Neo, Milford, NH, USA) of 2 mm-thick polymethylmethacrylate (PMMA) sheets. The depth of the culture chamber was 500 µm, while the foot height was 950 µm. Fabricated acrylic inserts were cleaned in an ultrasonic bath with soap water, then 40% ethanol solution, rinsed with water and blow-dried with air. Next, we used a double-sided adhesive tape (467-7952 MPL, 3M, Rüschlikon, Switzerland), which was laser-cut using a CO2 laser (Universal Laser System, Vienna, Austria), to attach the 0.4-µm-pore-size, hydrophilic PTFE biopore membrane (BGCM00010, Sigma-Aldrich, Buchs, Switzerland) to the insert. Before tissue culturing, inserts were UV sterilized in a cell culture hood for 20 min on each side.

### 3.2. Microfluidic setup for live-cell imaging

For tissue-slice culturing and live-cell imaging the LabSat instrument was modified to permit detachment of the imaging holder from the main unit. This allowed us to move the imaging holder into a laminar flow hood, enabling the loading of the insert from the 6-well culture plate to the imaging holder under aseptic conditions. We also equipped the imaging holder with a laser-cut, polycarbonate plastic support, which helped to align the tissue-culture insert (**Figure S1**). Additionally, the tubes delivering and retrieving the fluids from the imaging holder were cut to enable the detachment of the sample holder from the instrument for sample loading inside a cell-culture hood (**Figure S2**). Threaded fittings (XP-283, IDEX, Northbrook, Illinois) and unions (P-702-01, IDEX, Northbrook, Illinois) were used to connect/disconnect the tubes. To support tissue viability under perfusion, we included a gas exchanger, here called “oxygenator”, in the fluidic line upstream of the stainer, as described earlier.^9^ Briefly, the oxygenator consisted of a 50 ml Falcon tube (227261, Greiner Bio-One, St. Gallen, Switzerland) with a gas-permeable, 0.4-mm-ID PTFE tubing (S1810-06, Bola, Grünsfeld, Germany) coiled within the Falcon tube. We fully saturated the inside of the Falcon tube with carbogen gas (95% O2, 5% CO2). Re-equilibration with atmospheric gas partial pressures was avoided by using gas-impermeable PEEK tubing (150 µm ID, 1531, IDEX, Northbrook, Illinois) between the oxygenator and the imaging holder.

For live-cell imaging and culturing, the liquid flow rate was limited to 15-30 µl/min (**Figure S3**) by controlling the pressure differences between the main reservoirs and the outlet flask. Flow rates were assessed using an inline flow sensor (SLI-0430, Sensirion, Rüschlikon, Switzerland) downstream of the stainer.

Live-cell imaging experiments were performed on a Nikon Eclipse Ti2-E microscope (Nikon Europe B.V., Amsterdam, The Netherlands) equipped with an X-Light V3 spinning disk module (CrestOptics S.p.A, Rome, Italy). An environmental stage-top incubator (Life Imaging services, Basel, Switzerland) ensured that the system was kept at 37°C during imaging.

### 3.3. Cerebellar slice culture

Animal experimentation for mouse brain tissue harvesting was approved by the Basel cantonal Veterinary Office according to Swiss federal laws on animal welfare and was carried out in accordance with the approved guidelines.

Sagittal cerebellar brain slices were prepared as described previously.^45^ Briefly, the pups of wild-type C57BL/6JRj mice were sacrificed at P8–P11 by decapitation. Cerebella were harvested and placed in ice-cold Gey’s balanced salt solution (G9779, Sigma-Aldrich) containing 1 mM kynurenic acid (K3375, Sigma-Aldrich) and 6 mg/ml glucose (G8769, Sigma-Aldrich), further referred to as GBSSK. Cerebella were then embedded in sagittal orientation in 2% low-melting-point agarose (A9414, Sigma-Aldrich) prepared in GBSSK. The solidified agarose block was glued to the vibratome disc (VT 1200 S, Leica) using superglue. The disc was mounted in the vibratome chamber, filled with cold GBSSK, and 350-μm-thick slices were cut at a speed of 0.2 mm/s and an amplitude of 1.1 mm. After removal of excess agarose around the slices, the slices were transferred to Petri dishes filled with fresh, cold GBSSK on ice.

The cerebellar slices were then transferred onto either standard cell-culture inserts (PICM0RG50, Sigma-Aldrich) or onto the LabSat insert in 6-well plates with 1 ml of brain slice medium (BSM).^45^ We used fire-polished glass Pasteur pipets to place a maximum of three slices per standard tissue-culture insert or LabSat insert. The medium was exchanged daily, except for the day after transduction of the AAV, when the medium change was not performed. Slices were cultured in a humidified incubator with 5% CO2 at 37°C.

### 3.4. AAV transduction

Adeno-associated viruses (AAVs) were obtained from the viral vector facility of the University of Zurich. We used i) ssAAV-9/2-hGFAP-hHBbI/E-GCaMP6f-bGHp(A) at 2 × 10^9^ vg, ii) ssAAV-9/2-hGFAP-mCherry-WPRE-hGHp(A) and iii) ssAAV-PHP.eB/2-L7-6-jGCaMP8m-WPRE-bGHp(A) at 2 × 10^9^ vg to specifically label astrocytes or Purkinje cells in cerebellar slices. Viruses were diluted in PBS to achieve the indicated final viral titer concentrations in a volume of 5 µl PBS. On DIV4 and DIV6, 5 µl of the prepared virus solution were carefully pipetted on top of each slice during daily medium exchange. Slices were kept in the cell culture hood for an additional 10 min after transduction before returning them to the incubator for ALI culturing.

### 3.5. Calcium imaging

Calcium imaging assays were performed on a Nikon Ti-2 equipped with a spinning disc X-Light confocal unit as described before. After insert loading, slices were allowed to equilibrate to flow conditions for 3 flow periods, which consisted of 3 minutes flow and 9 minutes non-flow intervals, before imaging was started. We used a 40X water-immersion objective with a numerical aperture of 0.8. Conducted acquisitions in a single z-plane were 3 minutes long, using an exposure time of 300 ms with no binning in wide-field mode to acquire Purkinje cell calcium dynamics and 600 ms using 2×2 binning in the confocal mode for astrocyte calcium dynamics. All acquisitions were recorded during non-flow periods.

### 3.6. Glutamate stimulation

L-glutamic acid (G8415, Sigma-Aldrich, Darmstadt, Germany) was dissolved in BSM to a final concentration of 10 mM, and the pH was re-adjusted to 7.4 at room temperature. Next, the BSM containing L-glutamic acid was sterile-filtered with a 0.2 µm filter in a sterile hood and subsequently used to prepare the additional dilutions of 0.1 mM, 1 mM and 3 mM L-glutamic acid in BSM.

After replacing the normal medium with medium containing glutamate, slices were incubated for 10 min with the glutamate solution. Next, fresh glutamate was flown into the culture chamber three more times for a total stimulation duration of 30 min. Imaging of stimulated cells started 25 minutes after the glutamate containing medium entered the culture chamber.

### 3.7. Viability assay

Tissue health at the beginning and after 24 h of culture in the device, under the same pulsatile flow protocol of 3 min flow, alternated with 9 min no-flow periods, was assessed using cleaved Caspase-3/7 (Nucview 530, Biotium, Freemont CA, USA) and NucBlue (R37605, Thermo Fisher Scientific) to counterstain the nuclei of tissue slices. Imaging occurred 75 min after the staining solution entered the culture chamber. As a real-time control, we induced apoptosis after overnight culture by perfusing the slices with 40 µM Doxorubicin in BSM and incubating for 2 h before imaging.

### 3.8. Immunocytochemistry

Tissues were fixed for 1h in 4% PFA (28908, Thermo Fisher Scientific), diluted in 1X PBS on a shaker. After fixation, samples were cut out from the standard insert or the LabSat insertand stored in PBS at 4°C until staining. Tissues were permeabilized for 5 min in 0.25% Trypsin-EDTA (25200056, Thermo Fisher Scientific), followed by a 1 h blocking step in 1X PBS with 3% FBS, 3% BSA (diluted from A1595, Thermo Fisher Scientific) and 0.3% Triton-X100 (X100, Sigma-Aldrich). Primary antibodies were diluted in PBS with 3% FBS, 3% BSA, and samples were incubated overnight under agitation at 4°C. The samples were then washed three times for 10 min using PBS with 5% BSA. Secondary antibodies were diluted with DAPI (75004, Stemcell Technologies, Basel, Switzerland) in PBS containing 3% FBS and 3% BSA and incubated at room temperature for 3 h. Next, the samples were washed three times for 10 min with 1X PBS before mounting on 75 mm x 25 mm glass slides using ProLong Glass antifade mounting (P36984, Thermo Fisher Scientific) medium and covered with a #1.5 coverslip. The mounted samples were imaged with a Nikon Ti2 spinning disc confocal using a 10X dry and 40X water immersion objective.

The following primary antibodies were used: rabbit anti-Calbindin (1:1000 dilution, ab108404, abcam), chicken anti-Calbindin (1:1000 dilution, 214 009, Synaptic Systems), guinea pig anti-GFAP (1:2000 dilution, 173 308, Synaptic Systems), chicken anti-NeuN (1:500 dilution, ABN91, Sigma-Aldrich), and mouse anti-NeuN (1:1000 dilution, ab104224, abcam). As secondary antibodies, we used Alexa Fluor donkey anti-rabbit 647 (A21206, Thermo Fisher Scientific), Alexa Fluor donkey anti-chicken 488 (A78952, Thermo Fisher Scientific), Alexa Fluor donkey anti-mouse 405 (A48257, Thermo Fischer), and Alexa Fluor goat anti-guinea pig 555 (A21435, Thermo Fisher Scientific) at 1:200 dilution.

### 3.8. Image analysis

#### Tissue integrity

Purkinje cell somas were segmented into stacks of 10 µm, acquired at 0.4 µm z-step size, using Imaris v10.0.1 to determine the volume of each Purkinje cell soma and the number of cells per ROI. To normalize the number of cells with respect to the area of the Purkinje cell layer (PCL), we created maximum intensity projections of the ROIs. Then, the PCL boundary boxes were drawn in ImageJ (2.14/1.54f) using the polygon tool and measured, and the number of segmented cells was divided by the measured PCL area.

#### Tissue viability

Quantification of the intensity of the cleaved caspsase-3/7 signal was conducted by drawing squares of 300 px * 300 px in the projections of the maximum intensity of the acquired stacks and measuring the mean fluorescence intensity (MFI). Squares were drawn in both regions of the image containing tissue and tissue free regions. The MFI of tissue-free regions was subtracted from the MFI featuring tissue to determine the absolute intensity of the cleaved caspase-3/7 stain.

#### Calcium imaging

##### Astrocytes

We used AQuA2^46^ to analyze calcium dynamics in transduced astrocytes. Preprocessing of the acquired files included the removal of the first 21 frames of each acquisition, since we observed a strong intensity decay within the whole ROI, likely due to activation of calcium channels by the laser light ^47^. Batch processing of all acquired files was performed using Matlab (R2024a, The Mathworks, Inc., US) after confirming the settings using four different acquisitions. The following parameters were used for 40X acquisitions: median filter 0.5, Gaussian filter radius 1.5, intensity threshold 4.0, minimum duration 3, minimum size (px) 15, minimum seed size 0.02, Z score 3.0, maximum dissimilarity 0.5, minimum source size 0.01, sensitivity level 8. After batch processing, the results of the detected events were further filtered to include only events with an event area between 10-1500 µm^2^ for 40X images and an event duration between 1.6 and 60 s.

##### Purkinje cells

Calcium dynamics in transduced Purkinje cells were analyzed using Fiji. First, we removed the first 30 frames to adjust for a strong reduction in fluorescence intensity. Next, the background signal was subtracted using a rolling-ball method, and the image threshold was set using the IsoData algorithm. Next, watershed was applied, and the “Analyze particles” function was used to create masks of the somas and processes. If necessary, thin processes were manually added as cell ROIs using the polygon tool. Fluorescence traces were then measured using the in-built multi-measure function.

To calculate ΔF/F0, we used a custom python code (Python v 3.12.3) written in the Jupyter notebook environment. We determined F0 as the mean value of the 50 lowest recorded values per trace. Next, a Savitzky-Golay filter was applied to the raw trace for smoothening and to improve the spike detection. To be accounted as a true spike, traces had to deviate more than three times the standard deviation of the noise level. We considered the noise level as the difference between the raw trace and the smoothened trace. To adjust for a strong global decrease in fluorescence intensity, we calculated the ΔF/F0 amplitude based on the ratio of local minima (prominence) to the peak maxima of the smoothened trace. Per cell ROI, the mean value of all detected spikes was then used for visualization and statistics.

#### Cellular morphology

Morphological features of Purkinje cells (somatic area, thickness of dendritic processes and number of dendritic branches) were all measured in the Nikon software on maximum intensity projections of acquired z-stacks. The area of the Purkinje cell soma was determined using the automated ROI detection tool to mask the soma. The thickness of processes was measured by drawing three point-to-point lines per process and taking the average of these three measurements. We used the numbering tool to count the number of branches and astrocytic processes.

Astrocyte somas were segmented in Imaris (v 10.0.2) using the “cells” module based on intensity thresholding of a 50-µm stack acquired with 2 µm step size. After batch processing, each file was double checked for quality control before exporting soma volume values and the mean fluorescence intensity per soma for data visualization.

### 3.9. Data visualization and statistical analysis

GraphPad Prism v 10.5.0 was used to visualize the data and perform statistical analysis. The data were presented as individual values and mean values +/- SD, unless otherwise indicated in the figure caption. We first removed outliers from the data sets before testing for normality, followed by either an unpaired one-sided t-test or an appropriate one-way ANOVA, as indicated in the figure captions. Per condition, we acquired data from 5 slices cultured on 2 independent chips with 2-3 ROIs per slice for all Purkinje cell calcium metrics and morphology, astrocyte morphology and calcium dynamics of the 0 mM and 1 mM conditions. Data sets of astrocytes exposed to 0.1 mM Glutamate stem from 3 slices where 3-4 ROIs per slice were acquired.

## Acknowledgements

This work was financially supported by Innosuisse – the Swiss Innovation Agency- – under grant 120.665 IP-LS. The authors acknowledge the Single Cell Facility (SCF) at D-BSSE of ETH Zürich for help and support.

## Author Contributions

J.B.P. and M.M.M. conceived and developed the design of the culture insert and experimental paradigm. J.B.P. and M.M.M. designed the experiments and performed the experiments with the support of E.F.. L.H. adapted the fluidic system for live-cell imaging. A.H. and M.C. provided resources, acquired funding and supervised the project with M.M.M.. J.B.P. analyzed the data and wrote the manuscript; A.H. and M.M.M. edited the manuscript. All authors read the manuscript and approved the final version.

## Data Availability Statement

Data is available upon reasonable request from the corresponding author.

