## Supplementary information for "Microfluidic System for On-Demand Imaging of Organotypic Slices"

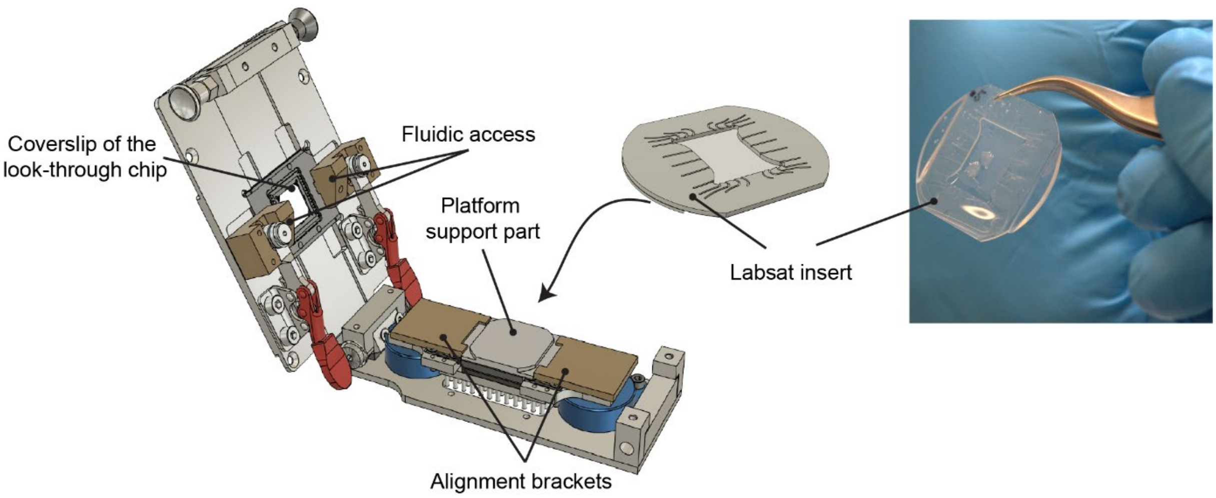


**Figure S1**: Details of the opened imaging holder, where the developed inserts were transferred to. The insert support part and the alignment brackets ensure a consistent placement of the insert in the imaging holder.

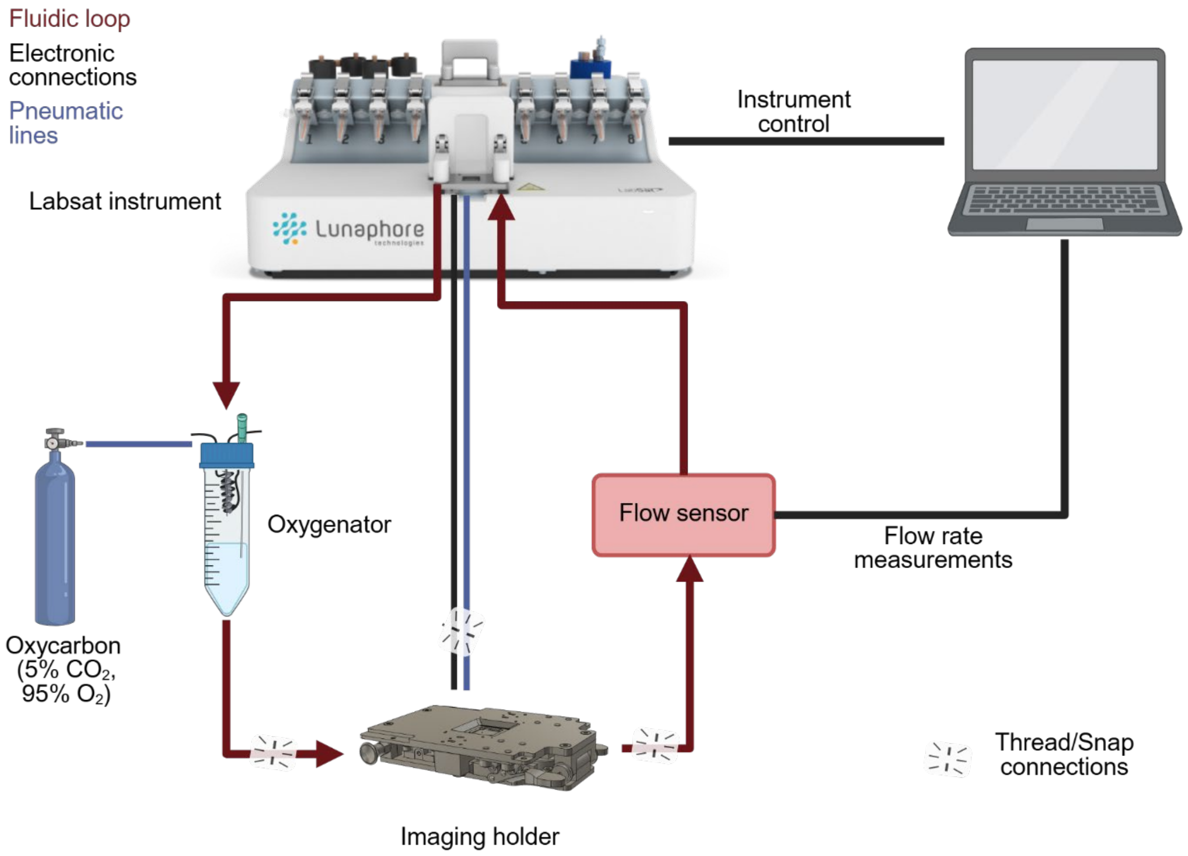


**Figure S2**: Schematic showing fluidic, electrical and pneumatic connections between the LabSat instrument and introduced adaptations to enable live-cell imaging.

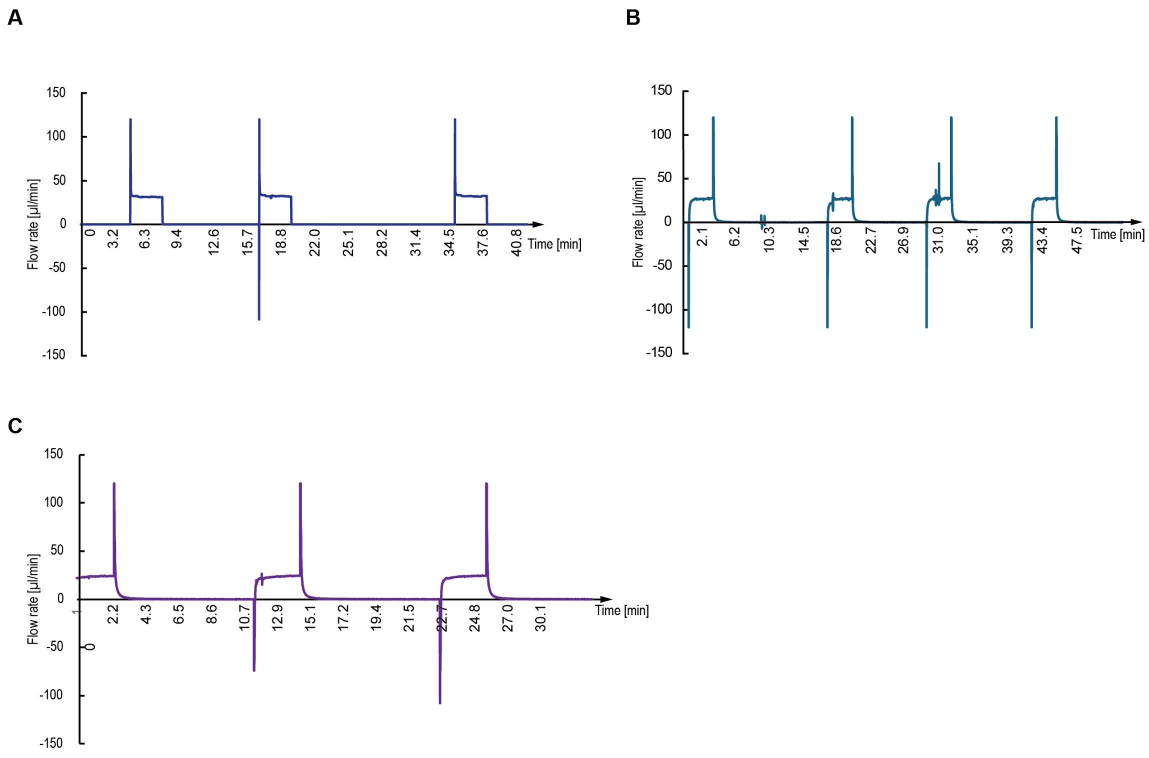


**Figure S3: A-C:** Exemplary flow traces of three different chips acquired through the flow sensor placed downstream of the imaging holder with the tissue slices.

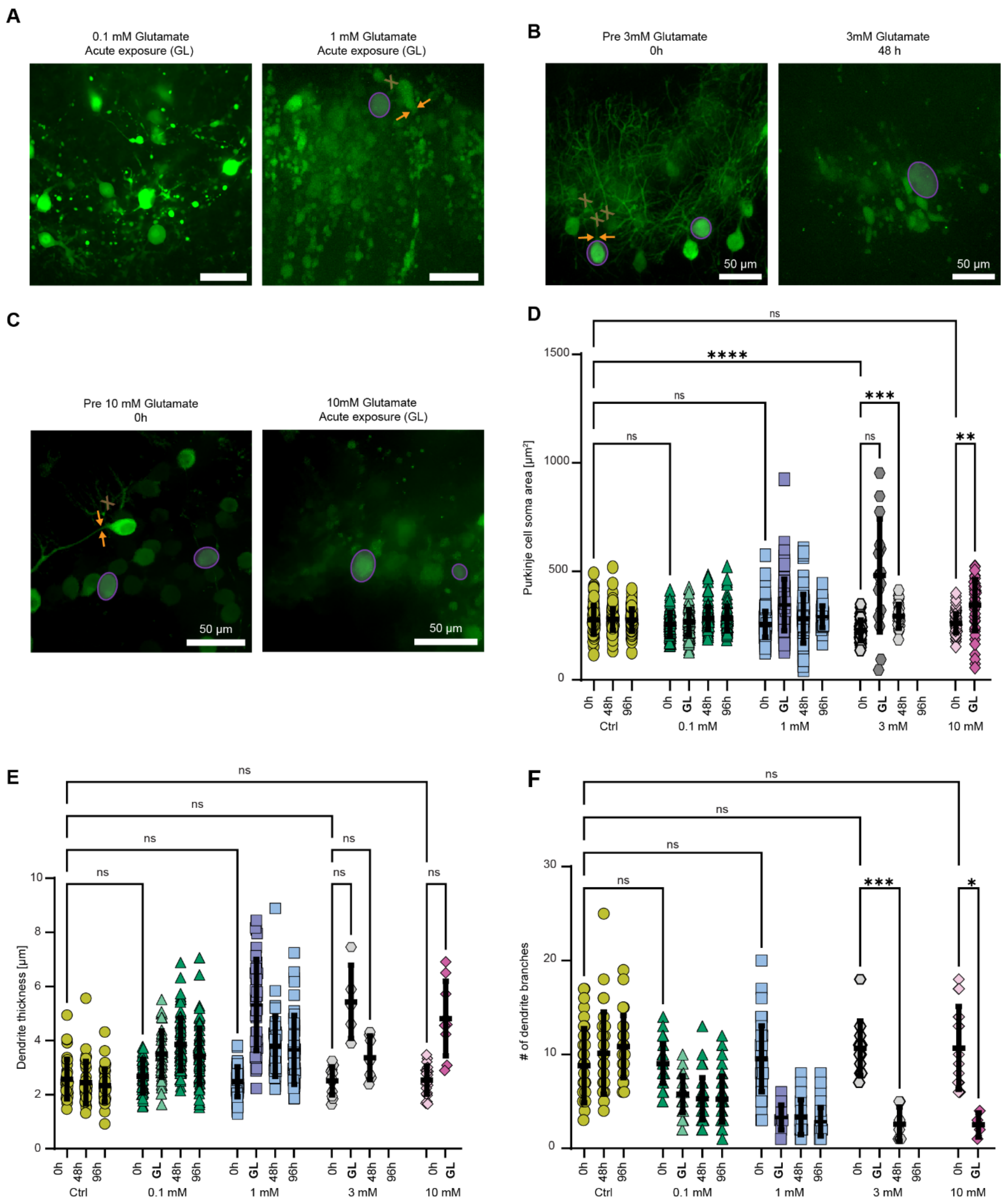


**Figure S4**: **A**: Maximum intensity projections of morphological features of Purkinje cells during exposure to 0.1 and 10 mM glutamate. **B-C**: Maximum intensity projections of morphological features of Purkinje cells prior to and during exposure to 3 mM glutamate and 10 mM glutamate. **D**: Purkinje cell somatic area measured at different time points before, during and after glutamate concentration. **E-F**: Dendrite thickness and number of dendritic branching points before glutamate exposure at 0 h, during glutamate exposure (GL), and 48 h and 96 h after glutamate exposure. Data from n=5 slices per condition for the control, 0.1 mM and 1 mM glutamate, n=2 slices for 3 mM and 10 mM glutamate exposure. Statistical analysis was performed using a one-way, non-parametric ANOVA test.
